# Epac activation reduces trans-endothelial migration of undifferentiated neuroblastoma cells and cellular differentiation with a CDK inhibitor further enhances Epac effect

**DOI:** 10.1101/2024.05.16.594493

**Authors:** Rabiu Inuwa, Diana Moss, John Quayle, Rasha Swadi

## Abstract

Neuroblastoma (NB) is the most common solid extracranial neoplasm found in children and is derived from primitive sympathoadrenal neural precursor. The disease accounts for 15% of all cancer deaths in children. The mortality rate is high in patients presenting with a metastatic tumour even with extensive treatments. This signifies the need for further research towards the development of new additional therapies that can combat not only tumour growth but metastasis, especially amongst the high-risk groups. During metastasis, primary tumour cells become migratory and travel towards a capillary within the tumour. They then degrade the matrix surrounding the pericytes and endothelial cells traversing the endothelial barrier twice to establish a secondary. This led to the hypothesis that modulation of the endothelial cell junctional stability could have an influence on tumour metastasis. To test this hypothesis, agents that modulate endothelial permeability on NB cell line migration and invasion were assessed *in vitro* in a tissue culture model. The cAMP agonist and its antagonists were found to have no obvious effect on both SK-N-BE2C and SK-N-AS migration, invasion and proliferation. Next, NB cells were cocultured with HDMEC cells and live cell imaging was used to assess the effect of an Epac agonist on trans-endothelial cell migration of NB cells. Epac1 agonist remarkably reduced the trans-endothelial migration of both SK-N-BE2C and SK-N-AS cells. These results demonstrate that an Epac1 agonist may perhaps serve as an adjuvant to currently existing therapies for the high-risk NB patients.

## Introduction

In children, neuroblastoma is the most common solid extracranial tumour, and the primitive sympathoadrenal neural precursor is the source of this tumour [1] Around 15% of paediatric cancer deaths are related to this condition [2]. Based on a combination of radiographic, histologic, cytogenetic, and age data at the time of diagnosis, patients with neuroblastoma are risk categorised [3].

Even with rigorous multimodal treatment including radiation therapy, immunotherapy, autologous stem cell transplantation, and the differentiation agent Retinoic acid (RA), patients with high-risk neuroblastoma have an overall survival rate of less than 50% [4,5]. The mortality rate is substantial for patients who present with a metastatic tumour. This highlights a need for novel treatments that not only restrict tumour growth but prevent metastasis, particularly in high-risk patients.

Primary cancer cells become migratory during metastasis and move into the blood vessels within the tumour (*intravasation*). After traversing the circulatory system, they break down the extracellular matrix enclosing the pericytes and endothelial cells, passing through the endothelial barrier (*extravasation*), to colonise and form a secondary tumour [6].

Many biological processes are known to be regulated by 3’,5**’-**cyclic adenosine monophosphate (cAMP), a product of the action of adenylate cyclase on ATP [7]. The compartmentalisation of proteins that generate, degrade, and activate the cAMP effectors in the cell provides evidence that the cAMP signalling pathways function in a stimulation-specific manner [8]. According to [9], cAMP enhances the endothelium barrier by activating exchange protein directly activated by cAMP (Epac1) within the vascular endothelial cells. This prompted us to hypothesise that modulation of endothelial cAMP and Epac 1 signalling may alter endothelial cell junctional stability, and therefore could have an influence on tumour metastasis.

In this study, in preliminary experiments we investigated the effect of agents that modulate endothelial permeability via Epac signalling on NB cell line migration and invasion *in vitro* by means of a tissue culture model. We demonstrated that neither an Epac1 agonist nor its antagonists affect the migration, invasion, or proliferation of SK-N-BE2C or SK-N-AS cells. Next, we examined co-cultured NB cells with Human Dermal Microvascular Endothelial Cells (HDMEC). Our findings demonstrate that an Epac1 agonist significantly inhibits the transendothelial migration of SK-N-AS and SK-N-BE2C cells by around 52%. Furthermore, we observed a two-fold increase in the percentage of retained cells for the Palbociclib differentiated (>90%) compared to the RA differentiated cells, suggesting a potential therapeutic mechanism for treating high-risk neuroblastoma patients.

## Materials and Methods

### Cell culture

#### Routine Culturing of Neuroblastoma cell lines

For neuroblastoma cell lines, the cells were cultured in serum-containing medium made of DMEM (Gibco), 10% FBS (Cat no. 10270106 Life Technologies) and 1% NEAA (Sigma, M7145). Cells were generally cultured as monolayers in either 25 cm^2^ or 75 cm^2^ tissue culture flasks (Corning, UK), using the appropriate media. Flasks were kept in a humidified incubator at 37°C, 5% CO_2_ and routinely passaged as they reached 80-90% confluency. The cells were routinely tested for mycoplasma to ensure they remained free of contamination. GFP-labelled cells and wild type cells were cultured in the same medium using the same methods [10,11].

#### Routine Culturing of HDMEC cells

Cryopreserved human dermal microvascular endothelial cells (HDMECs, Cat no. C 12221) were purchased from Promocell (Heidelberg, Germany) at passage 2 (P2). HDMECs were grown in 0.1% gelatin-coated 75 cm^2^ flasks with 12 ml fully supplemented Endothelial cell growth medium MV 2 kits (C-22022 PromoCell GmbH) in a 5% CO_2_ humidified incubator at a temperature of 37°C. Cells were routinely passaged when reaching about 80-90% confluency.

### Compounds

#### Activators (agonists)

These include the cyclic AMP elevating agent Forskolin (FSK) (CN-100-0010, Enzo Life Sciences Inc, USA) and Epac1 activator Epac non-acetomethyl ester (non-AM) 8-(4-Chlorophenylthio)-2’-O-methyladenosine-3’, 5’-cyclic monophosphate (8-pCPT) (C 052, Biolog, Life Science Institute, Germany).

#### Inhibitors (antagonists)

The inhibitors used include Epac inhibitors (ESI-09 and HJC-0197), protein kinase A (PKI) inhibitor. ESI-09 (B 133) and HJC-0197 (C 136) were bought from Biolog, Life Science Institute, Germany. PKI (P 210) (myristoylated PK (14-22)) amide) was bought from Enzo Life Sciences Inc, USA.

#### Differentiation agents

Agents used include All-Trans retinoic acid (RA) (CAS no. 302-79-4 Sigma-Aldrich) and Palbociclib (Cat no. S1579, Selleckchem, US).

### Transendothelial migration assay

Transendothelial migration study was carried out firstly by generating a 100% confluent HDMEC monolayer and then subsequently co-culturing with the GFP labelled NB cells. Sterile size 1 coverslips (diameter, 13 mm) were placed per well of a 24 well plate and then coated with 0.1% gelatin (Cat no. 39465, Sigma) for 10-15 minutes in a 37°C, 5% CO_2_ incubator. After incubation, the gelatin solution was aspirated and discarded. Cells were trypsinised and counted with a haemocytometer. HDMECs were seeded onto coverslips at a density of 1 x 10^5^ cells per well in 500 µl fully supplemented endothelial cell culture medium (Promocell) to generate a 100% confluent monolayer. Endothelial monolayer cells were then incubated with 500 µl of pre-warmed 1:1 medium for 30 minutes at 37°C, 5% CO_2_ for equilibration. The equilibration medium was then aspirated and 3 x10^4^ GFP labelled NB cells were gently seeded onto the endothelial monolayer in 500 µl (1:1) medium with or without treatment compounds. The culture plate was mounted on Cell IQ machine (CM Technologies, Finland). 4-5 positions were selected per well and serial images for two channels (Green and Gray) were captured every 30 minutes for 24 hours. The analysis was done by manual counting of the green cells on top of the endothelial monolayer (grey) at 12 hours.

### Immunocytochemistry

For HDMECs, 13 mm coverslips were coated with 0.1% gelatin, placed in a 24-well culture plate, and incubated for 10-15 minutes in a 37°C / 5% CO_2_ incubator. After incubation, gelatin was carefully aspirated and discarded from each well. HDMECs were seeded onto coverslips with a density of 1 x 10^5^ cells per well in 500 µl of fully supplemented cell culture medium (Promocell), and 3 x 10^5^ cells per glass bottom surface of a 35 mm glass-bottom dish (Greiner) in 1.5 ml of fully supplemented cell culture medium (Promocell). Cells were cultured at 37°C / 5% CO_2,_ typically for 24 hours. Culture medium was removed, and cells were washed with 1 ml DPBS. Cells were fixed with 4% paraformaldehyde for 10 minutes and replaced with 500 µl 100 mM glycine solution (pH 7.4) for 10 minutes at room temperature. Cells were incubated in permeabilising solution and then washed with PBS, 3 times for 5 minutes. Coverslips were then incubated in blocking solution for 30 minutes and then incubated with 100 μl antibody diluting solution containing 2% v/v goat serum, 0.05% v/v Triton X-100, 1% w/v bovine serum albumin in sodium citrate buffer (SSC, containing, in mM; 150 NaCl and 15 Na_3_ Citrate), pH 7.2 for 30 minutes at room temperature. 70 μl of 1:100 Anti-human VE-cadherin primary antibody (Monoclonal, Mouse IgG) was added to the wells and incubated at 4°C overnight. The secondary antibody (Alexa Fluor 488-tagged goat anti-mouse secondary antibody used at 1:500 and DAPI were diluted in blocking solution, applied to each coverslip, and incubated in the dark at room temperature for 1 hour. The cells were then washed with antibody diluting solution three times at room temperature, 10 minutes each. Coverslips were washed quickly with UHQ water twice and left to air dry in the dark. When dry, coverslips were mounted onto glass slides with mounting medium (Dako, S3023). Finally, the slides were stored at 4°C to be visualised.

### Confocal microscopy

An LSM800 confocal microscope was used to visualise the labelled cells with a 63x oil immersion objective. Alexa Fluor 488 was excited at 488 nm using an argon-ion laser, and emission light was collected at 500-550 nm. For the detection of DAPI nuclear staining, the working excitation was 405 nm. mPlum-VE-cadherin-N-10 was excited at 590 nm.

### Confocal microscopy (live cell imaging)

LSM800 confocal microscope was used to collect fluorescence signal during the live-cell imaging study in controlled humidity and temperature of 37°C / 5% CO_2_. Data from were analysed with Zen SP1 software (Carl Zeiss AG, Oberkochen, Germany).

### Statistical Data analysis

All data obtained were first expressed as a mean with the standard error of the mean (SEM), unless otherwise stated. Statistical analyses were performed using GraphPad Prism version 6.00. Statistical comparison of the mean of two datasets was done using paired / unpaired Student’s t-test. One-way analysis of variance (ANOVA) was used for the comparison of the mean of more than two data sets, followed with Tukey’s post hoc analysis. P <0.05 was considered statistically significant. Statistical significance indicated as * P < 0.05, ** P<0.01 and *** P < 0.001.

## Results

### Co-culturing model: Optimisation of HDMEC and NB cells (transendothelial migration model)

Previous work had identified a potential role of the Epac pathway in strengthening endothelial adherens junction [12]. To further clarify the effect of the Epac activator 8-pCPT on human dermal microvascular endothelial cells (HDMEC), and to investigate its role in the process of trans-endothelial migration of NB cells, it became necessary to establish a model to study the transmigration of NB SK-N-BE2C and SKNAS cells across the endothelium. We developed an improved co-culture model for HDMECs and NB cells for trans-endothelial migration study. Most previous trans-endothelial studies were carried out using transwells [13,14], which have drawbacks because it is difficult to visualise the inner aspect of the transwell membrane. In addition, endothelial cells are difficult to culture as a confluent monolayer, and any disruption to the monolayer will invariably affect the real outcome of the study.

To overcome these issues, we coated glass coverslips with 0.1% gelatin then seeded 1.0 x 10^5^ HDMECs and cultured for 24 hours using a 24 well plate. At the end of 24 hours, GFP-labelled NB cells were seeded in (1:1) endothelial cell medium: NB cell culture medium. The plate was then mounted on the cell IQ and serial imaging was carried out for 24 hours (**Fig 1**).

**Fig 1.**
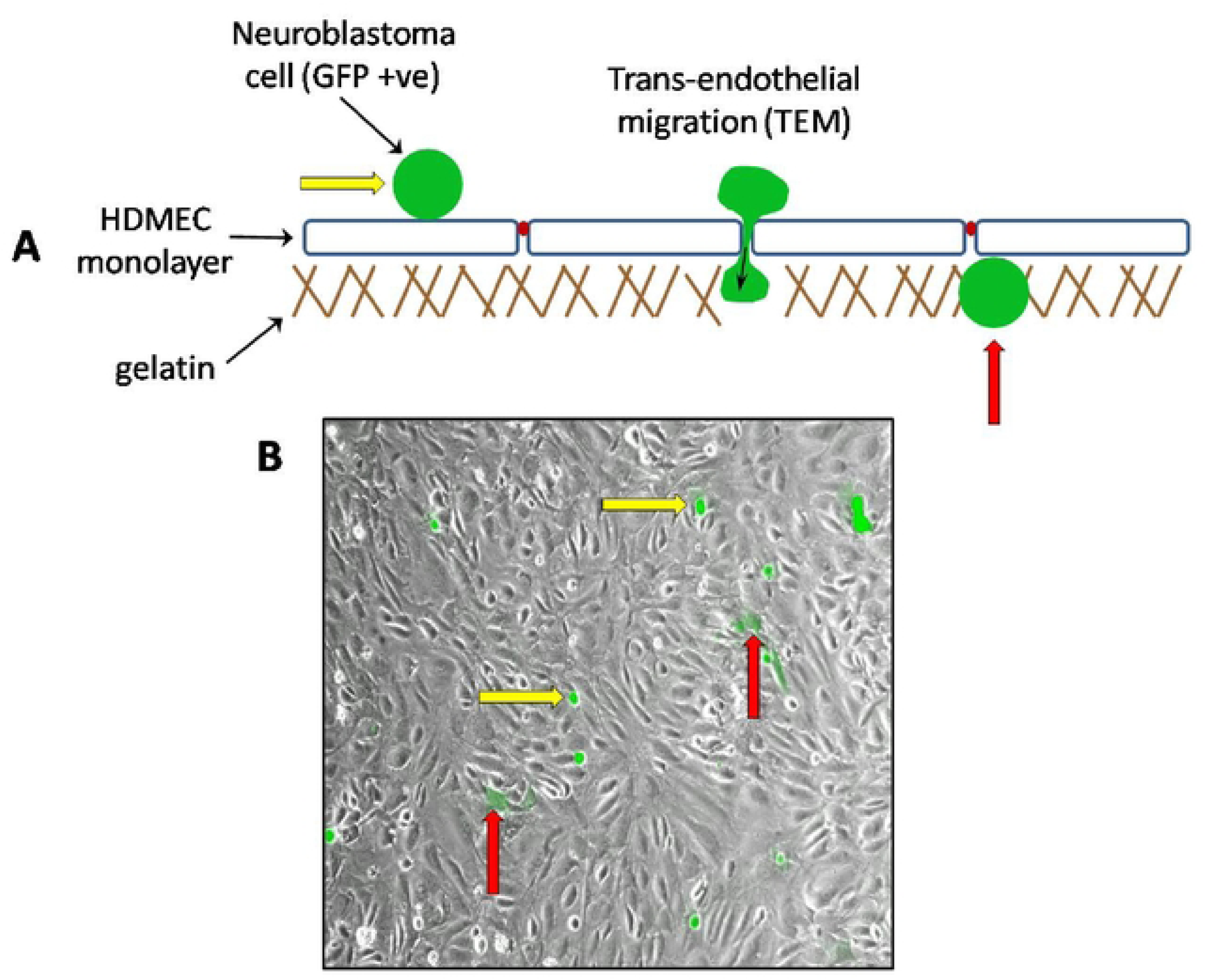
Trans-endothelial migration model. The illustration (**A**) above shows a diagrammatic representation of the trans-endothelial migration model. HDMECs (1 x 10^5^ cells) were seeded on a glass coverslip and incubated for 24 hours. GFP-labelled NB cells, as indicated by the yellow arrows (**A** & **B**), were then seeded on top of the HDMEC monolayer (grey). Transmigrated cells are seen to fade in colour as they traverse the endothelial cells monolayer as shown by the red arrows (**A** & **B**). Image was captured using cell IQ machine.

### Epac agonist strengthens the HDMEC adherens junctions

HDMECs were cultured for 24 hours and then treated with 8-pCPT for 30 minutes. Both treated and untreated cells were stained with mouse anti-VE-cadherin, in order to clearly distinguish the effect of treatment on the HDMECs adherens junctions. As expected, within 30 minutes of treatment with 8-pCPT, the HDMECs exhibited a re-distribution of the VE-cadherin around the cell membranes (**Fig 2**), as also reported in other studies [15–18]. This is indicative of the strengthening of the VE-cadherin adherens junction compared to the control. The effect of Epac1 agonist treatment is most obvious in the VE-cadherin and DAPI merged images captured using a confocal microscope (**Fig 2**).

**Fig 2.**
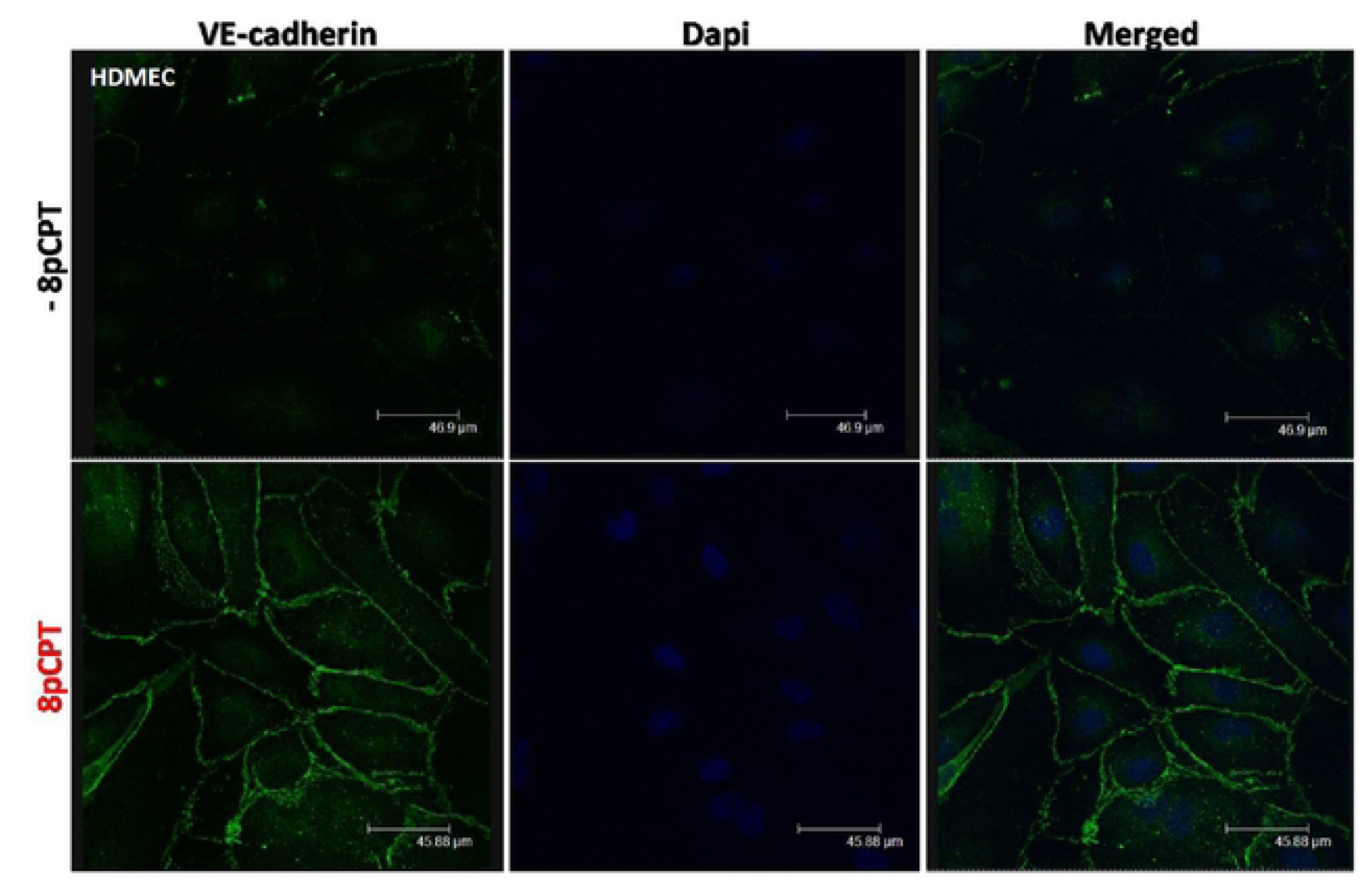
Immunocytochemistry of VE-cadherin expression in HDMECs following 8-pCPT treatment. VE-cadherin distribution in HDMECs was primarily identified as discontinuous membranous sections. Following the treatment with 10 µM 8-pCPT for 30 minutes at 37°C & 5% CO_2_ the treated cells demonstrated a well-formed continuous VE-cadherin border along the membrane of adjacent HDMECs. The nuclei are stained with DAPI (blue). Images were captured with a confocal microscope. This experiment was repeated twice (N=2).

### Characterisation of trans-endothelial cellular migration of SK-N-BE2C and SK-N-AS

Next, we characterised the transmigration of cells from two NB cell lines, SK-N-BE2C and SK-N-AS cells, across the HDMEC monolayer. 3 x 10^4^ GFP-labelled NB cells were co-cultured with HDMECs, as described above (**Fig 1**). Co-cultured cells were mounted on cell IQ for 24 hours. At least four different fields containing GFP-expressing cells were selected and imaged serially every 30 minutes. To quantify the transmigration of NB cells across the endothelium, GFP-labelled NB cells on top of the endothelial cells were counted at 60 minutes after seeding and also at the end of 12 hours. Data are expressed as percentage of non-migrated cells at 12 hours versus the total cells at 60 minutes (**Fig 3**). The majority of cells from both cell lines migrated through the endothelial monolayer (81% SK-N-AS and 82.7% SK-N-BE2C). SK-N-AS cells have a slightly greater percentage of non-migrated cells relative to SK-N-BE2C cells at 12 hours, although the observed difference (1.75%) was not statistically significant (**Fig 3**).

**Fig 3.**
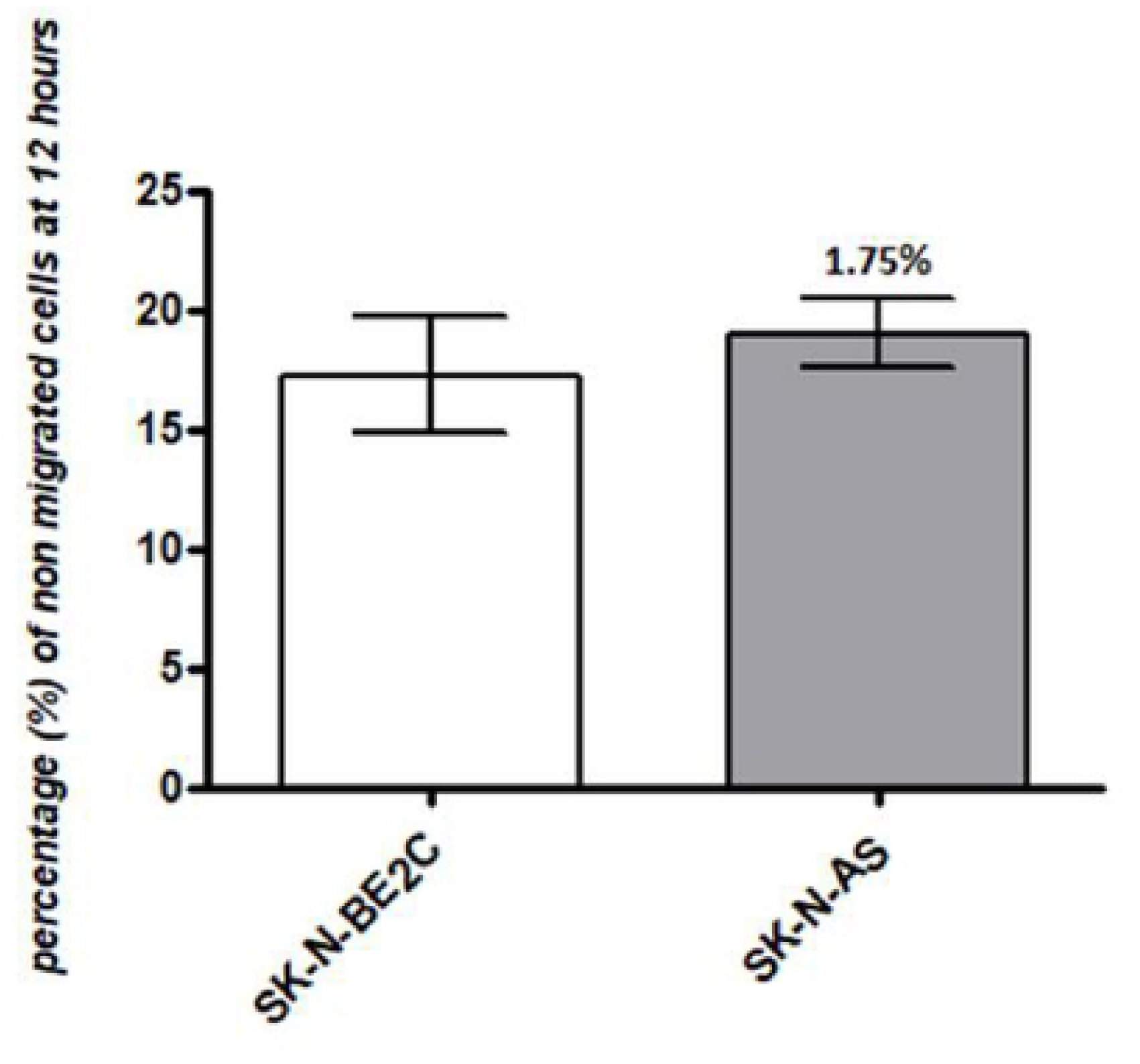
Comparison of trans-endothelial cell migration between SK-N-BE2C and SK-N-AS cells. For each of the two NB cell lines, 3 x 10^4^ GFP-labelled cells were seeded on a 100% confluent HDMEC monolayer. Serial images were captured at 30 minutes intervals using cell IQ machine. Percentage of non-migrated NB cells were quantified by counting the GFP cells at 60 minutes and at the end of 12 hours. Three separate independent experiments were analysed for each of the NB cell lines. The Figure shows that SK-N-BE2C cells have a smaller percentage of non-migrated cells compared to SK-N-AS cells (1.75%) The percentage of migrated **SK-N-BE2C** cells is 82.7%, while for **SK-N-AS** is 80.9%. Each bar represents the mean ± SEM. Students t-test was used to compare the percentage of non-migrated cell in the two cell lines. The difference was not statistically significant (P > 0.05).

### Epac agonist (8-pCPT) reduces trans-endothelial cellular migration of SK-N-BE2C and SK-N-AS cells

Previously, we demonstrated that agonists and antagonists of PKA and Epac do not have any observable effects on NB cell line migration, invasion or proliferation (**See supporting information**). Our preliminary treatment of HDMECs with 8-pCPT for 30 minutes showed evidence of strengthening of VE-cadherin adherens junctions **Fig 2**, as reported in the literature [16–19]. As such we wanted to examine the role of activation of the Epac pathway on strengthening the endothelial cell junctions vis a vis transmigration of the NB cells across the endothelial barrier. GFP-labelled NB cells were plated on a confluent HDMEC monolayer with and without 8-pCPT treatment for 24 hours. Percentages of non-migrated GFP-labelled NB cells were determined at the end of 12 hours incubation relative to 1 hour post-plating the cells.

Treatment with 8-pCPT (10 µM) was shown to increase the percentage of retained (non-migrated) cells across the endothelium from 25.3% to 38.4% for SK-N-BE2C cells and from 28.1% to 44.1% for SK-N-AS cells, which is by about ∼52% in the case of both SK-N-BE2C and SK-N-AS cells relative to the control (**Fig 4A** and **B**). Furthermore, the time course analysis shows that at every point of analysis (1, 2, 6 and 12 hours) more cells were retained on top of the endothelial cells in treated condition compared to the untreated cells (**Fig 4C** and **D**), indicating that the effect of 8-pCPT treatment on HDMECs seems to be sustained during the course of treatment. The effectiveness of 8-pCPT was further evident by a higher area under the curve relative to the control as shown (**Fig 4E** and F).

**Fig 4.**
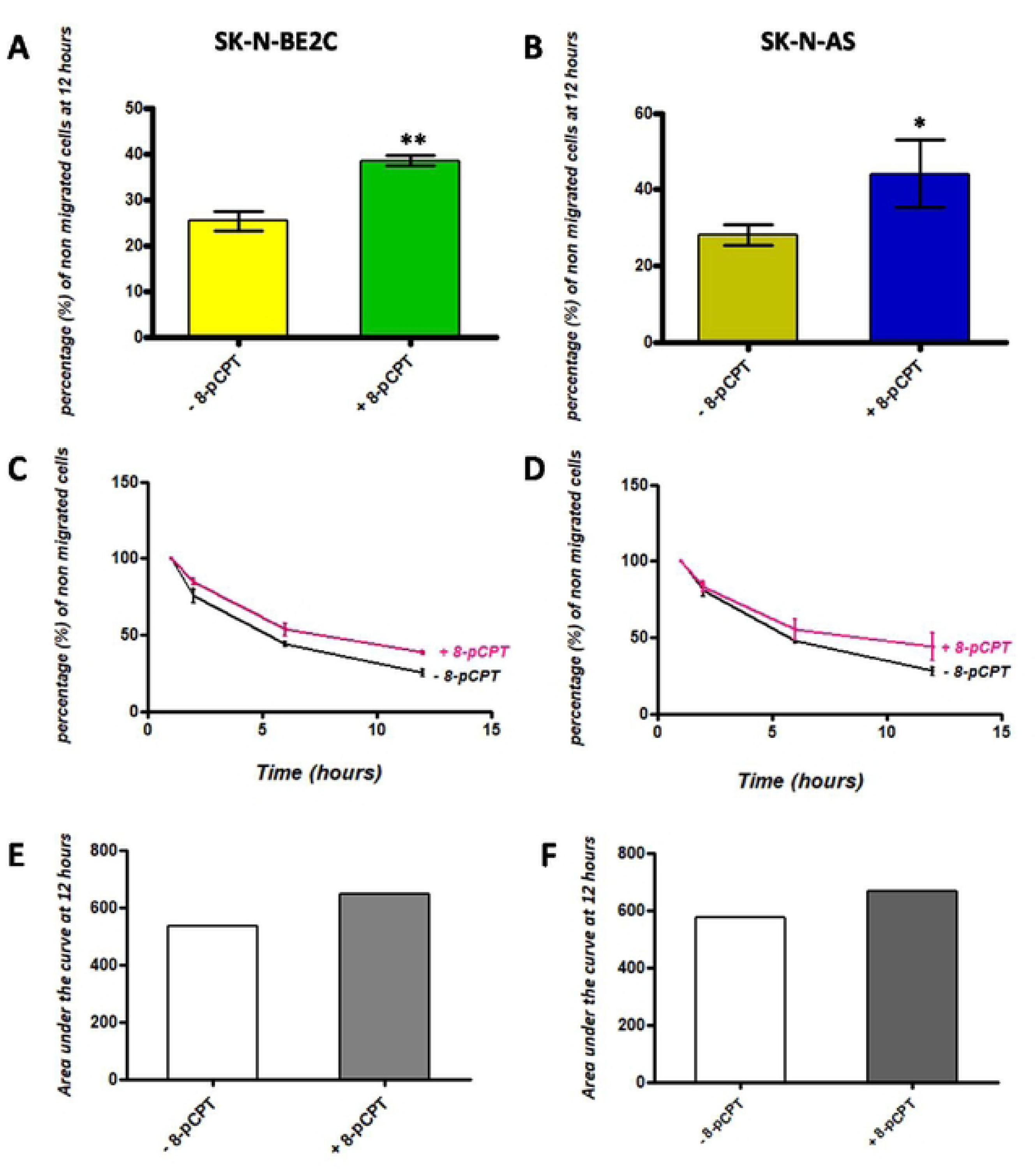
Effect of 8-pCPT treatment on trans-endothelial cell migration of SK-N-BE2C and SK-N-AS cells. For each of the two NB cell lines, 3 x 10^4^ GFP-labelled cells were seeded on a 100% confluent HDMEC monolayer with 8-pCPT (10 µM). Serial images were captured at 30 minute intervals using a cell IQ machine. Percentages of non-migrated NB cells were quantified by counting the GFP-labelled cells at 60 minutes and at the end of 12 hours. Three separate independent experiments were analysed for each of the NB cell lines and 12 fields per experiment. Each bar in **A** & **B** represents the mean ± SEM. **C** & **D** shows the time course effect following 8-pCPT treatment**. E & F** demonstrate higher area under the curve following 8-pCPT treatment at the end of 12 hours. Student’s t-test was used to compare the control (-8-pCPT) and treatment (+ 8-pCPT) conditions. There was a significant statistical difference between the 8-pCPT treatment and control for both SK-N-BE2C and SK-N-AS cells. P < 0.05 (*), P < 0.01 (**).

### HJC0197 or ESI-09 do not increase the trans-endothelial cellular migration of SK-N-BE2C and SK-N-AS cells

Since 8-pCPT is an agonist of Epac1 and we have shown it to lead to a significant decrease in the number of transmigrated cells (**Fig 4**), it was important to test the effect of Epac inhibitors on 8-pCPT augmentation of the trans-endothelial migration of NB cells. Epac inhibitors ESI-09 and HJC0197 were tested individually on the trans-endothelial migration of SK-N-BE2C and SK-N-AS cells. HJC0197 was reported to block both Epac1 and Epac2 mediated activity in HEK293T cells [20,21]. ESI-09 also has been shown to inhibit the activity of Epac1 and Epac2 and was previously reported to suppress the cell migration and invasion in pancreatic cancer cells [22–24]. ESI-09 and HJC0197 were used at 5µM, a dosage which falls around the IC_50_ of HJC0197 (5.9µM) and slightly above the reported IC_50_ of ESI-09 (1.4-3.2µM).

We observed no difference in trans-endothelial cellular migration between the control and the presence of inhibitors (P > 0.05) (**Fig 5A, B, C** and D). As previously, 8-pCPT increased the percentage of non-migrated cells across the endothelium compared to control (**Fig 5A** and B). Time course analysis shows that at every point of analysis (1, 2, 6 and 12 hours) more cells were retained on top of the endothelial cells in 8-pCPT treated condition compared to the HJC0197 and ESI-09 treated cells (**Fig 5C** and D). Further analysis of the area under the curve of **Fig 5C** and **D** showed a higher area for the agonist compared to the antagonist during the course of 12 hours treatment (**Fig 5E** and **F**).

**Fig 5.**
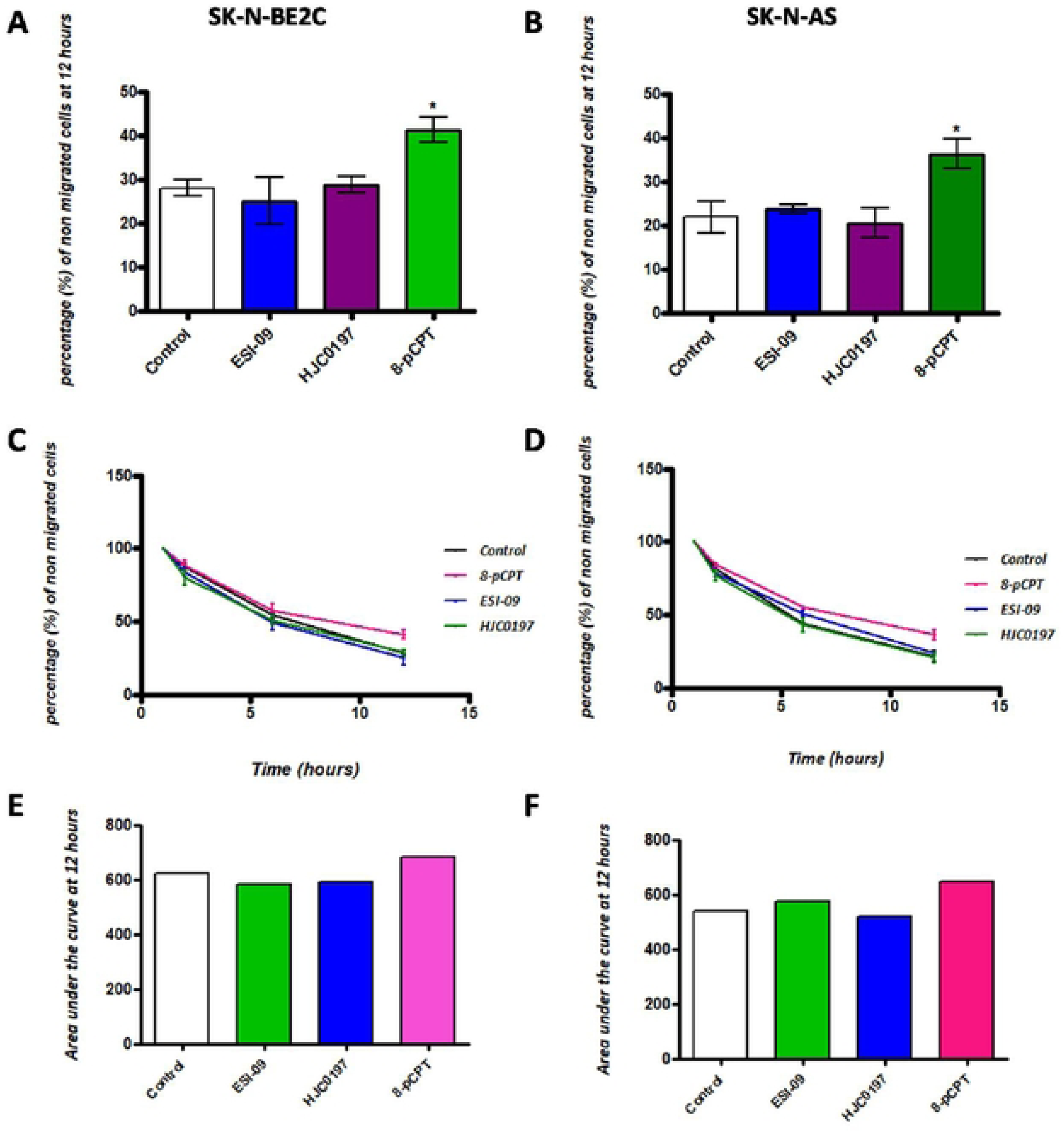
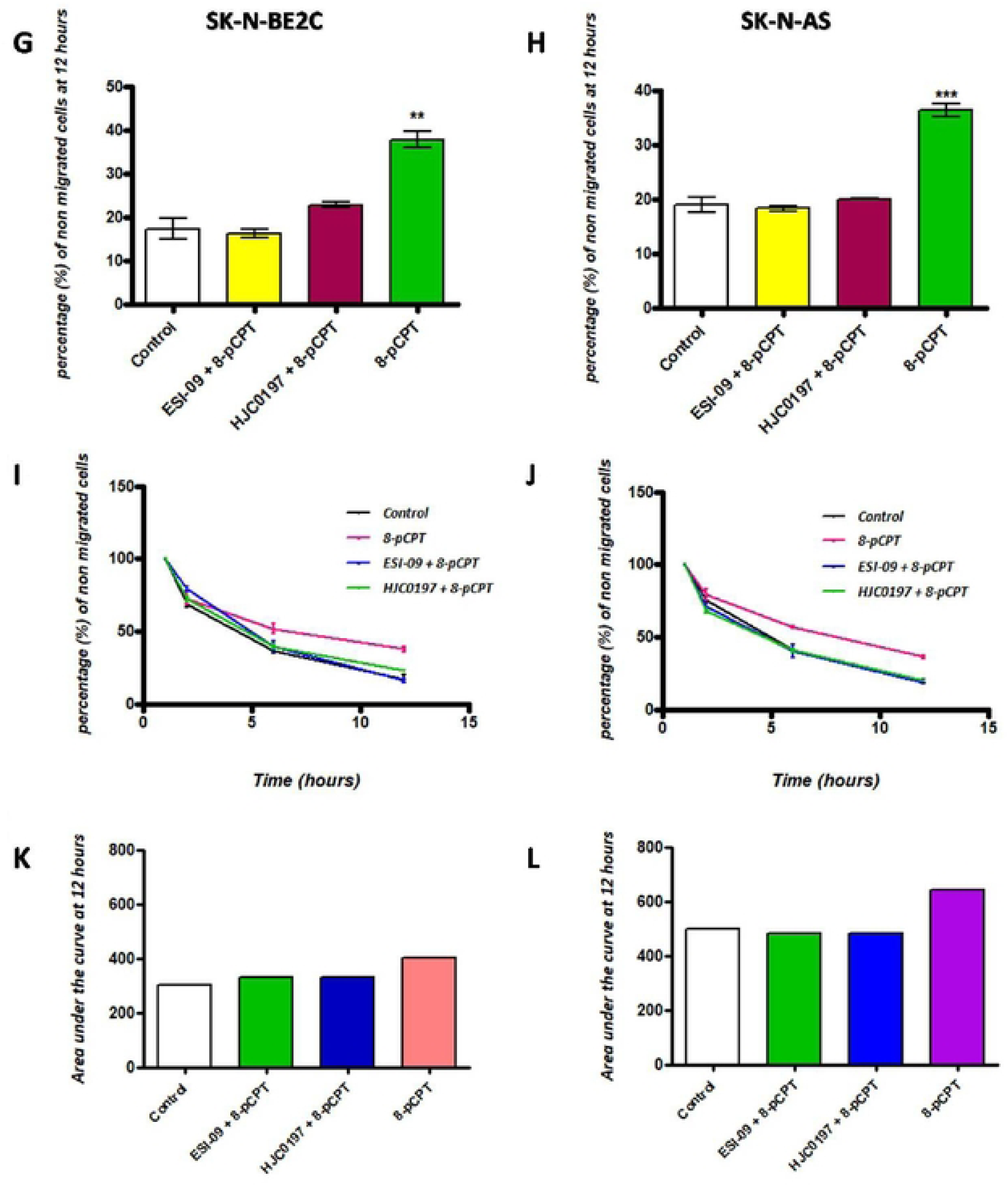
Effect of treatment with Epac agonist in combination with antagonist on trans-endothelial cell migration of SK-N-BE2C and SK-N-AS cells. HDMECs were plated on a 0.1% gelatin-coated coverslip in a well of a 24 well plate for 24 hours to generate a confluent monolayer. Serial images were captured at 30 minutes intervals using a cell IQ machine. Percentage of non-migrated NB cells were quantified by counting the GFP positive cells at 60 minutes and at the end of 12 hours. Three separate independent experiments were analysed for each of the NB cell lines. Each bar in **A**, **B, G** & **H** represents the mean ± SEM. **C**, **D, I** & **J** shows the time course effect following 8-pCPT treatment**. E, F, J & K** demonstrate higher area under the curve following 8-pCPT treatment at the end of 12 hours. One way ANOVA was used to compare the effect of treatment on NB cell trans-endothelial migration. Means comparison was assessed by Tukey post-test. P < 0.05 (*), P < 0.01 (**) and P <0 .001 (***).

### Epac antagonists prevent the effect of 8-pCPT on trans endothelial migration of SK-N-BE2C and SK-N-AS cells

Having shown that there is no significant difference in the pattern of trans-endothelial migration in the presence of two Epac antagonists when compared to control, we proceeded to investigate whether the antagonists are able to prevent the effect of Epac agonist 8-pCPT, to ensure the reduction in transmigration due to the agonist is not because of an off-target effect.

As previously, live cell imaging was performed using the cell IQ and transmigration was assessed during the 12 hours course of treatment. The percentage of the non-migrated SK-N-BE2C and SK-N-AS cells in the combination treatment (i.e. agonist + antagonist) condition was not significantly different from the control (P > 0.05, **Fig 5G** and **H**). This suggests that inhibition of the activity 8-pCPT by ESI-09 or HJC0197 prevents the effects on transmigration [20–24]. Further analysis of the area under the curve of **Fig 5I** and **J** showed a larger area for the agonist compared to both combination and control conditions during the course of 12 hours treatment (**Fig 5K** and **L**).

### Effect of cell differentiation on trans-endothelial migration of NB cells

Strengthening of the VE-cadherin mediated adherens junctions following treatment with 8-pCPT reduces the transendothelial cellular migration of both SK-N-BE2C and SK-N-AS cells, while combination treatment with Epac agonist and antagonist demonstrate a similar pattern of trans endothelial NB cellular migration to the control. We next wanted to examine any potential role of cellular differentiation of NB cells on transmigration across the endothelium. The question was whether cellular differentiation will confer any advantage in further reducing the trans endothelial migration of NB cells.

Various agents are used to differentiate NB cells. However not all NB cell lines are found to be sensitive to these agents. As such, in order to differentiate the NB cells, we used two known chemotherapeutic agents, retinoic acid (RA) and Palbociclib (a CDK4/6 inhibitor). Retinoic acid is one of the conventional agents often used in clinics for the treatment of high-risk cases of NB [25,26]. Palbociclib has been reported to induce cellular differentiation in breast cancer [27–30]. In addition, it has been used in our laboratory for the differentiation of various NB cell lines [10,11]. Amongst our two selected NB cell lines only SK-N-BE2C cells have been reported in our laboratory to be sensitive to both RA and Palbociclib treatment.

### Cellular differentiation of SK-N-BE2C cells using RA

We adopted the dosage used in the literature for RA, at a final concentration of 10 µM (31,32). SK-N-BE2C cells were plated on a glass coverslip in a 24 well plate. Then, attached cells were treated for 72 hours with RA at 5% CO_2_ and 37°C. Nuclei were stained with DAPI and images were captured. RA induced changes in the morphology of the cells (**Fig 6**) as indicated by neurite formation in most of the cells. Furthermore, at the end of 72 hours treatment, we passaged the treated cells and re-plated them in ordinary culture medium to further assess whether the effect of treatment is transient or permanent, prior to co-culturing with HDMECs.

**Fig 6.**
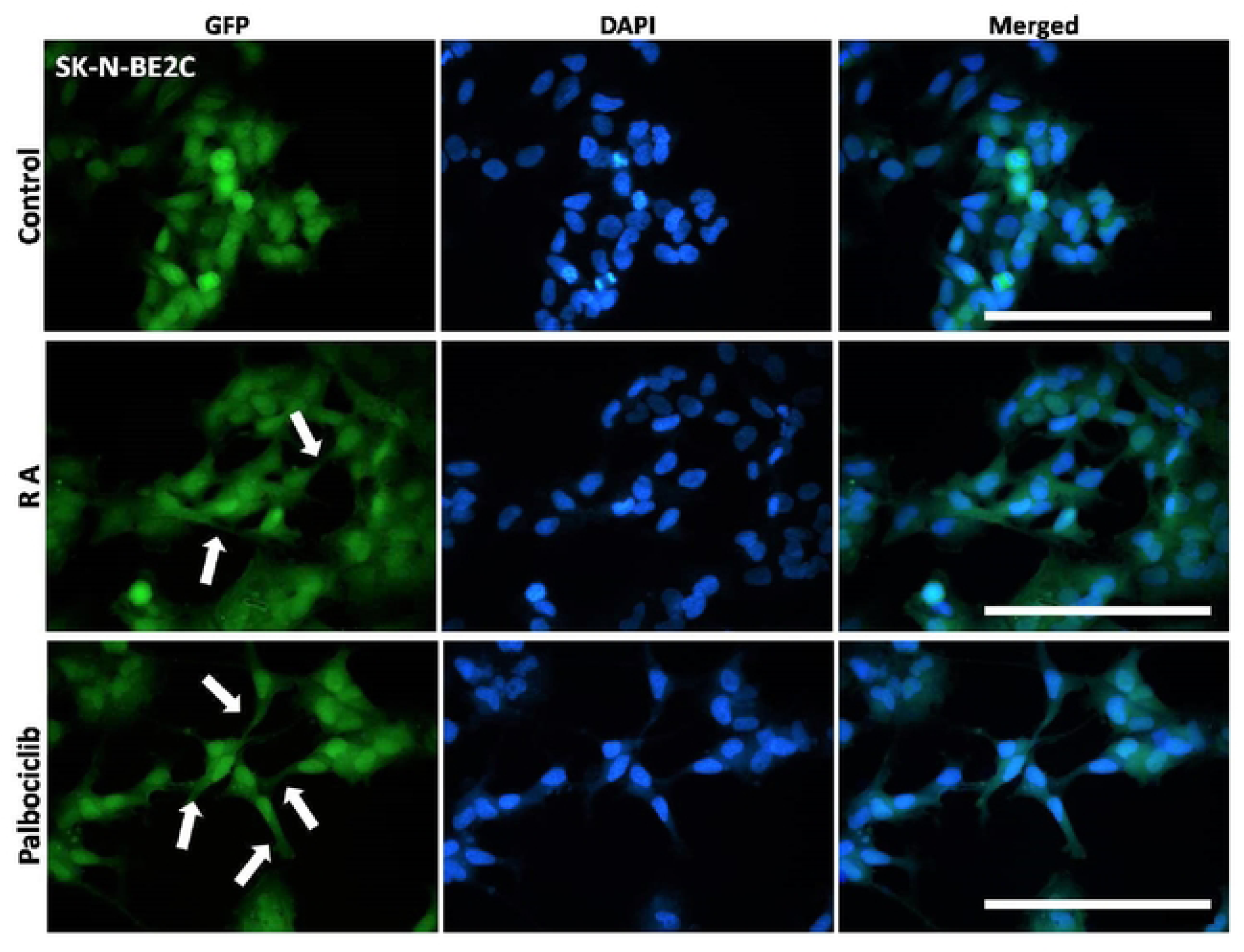
Morphological effect of RA and Palbociclib treatment on SK-N-BE2C cell morphology. SK-N-BE2C cells were cultured in a 24 well plate and treated with DMSO, RA and Palbociclib. GFP-labelled SK-N-BE2C cells are seen in green and DAPI stained nuclei in blue. SK-N-BE2C cells were treated for 72 hours (3 days) with 10 µM RA or with 5 µM Palbociclib. DMSO was used as control. RA acid treated cells show neurite formation (indicated by white arrows), which are more numerous in Palbociclib treated cells. DMSO treated cells maintain a round, undifferentiated form. Images were captured with a x40 objective. Scale bar is 250 µm. This experiment was repeated three times (N=3).

Following replating in a plain culture medium, we observed that the cells continued to form processes (**Fig 7A**) up to the time they become 100% confluent, and also even after another passaging, indicating that the effect of treatment is sustained.

**Fig 7.**
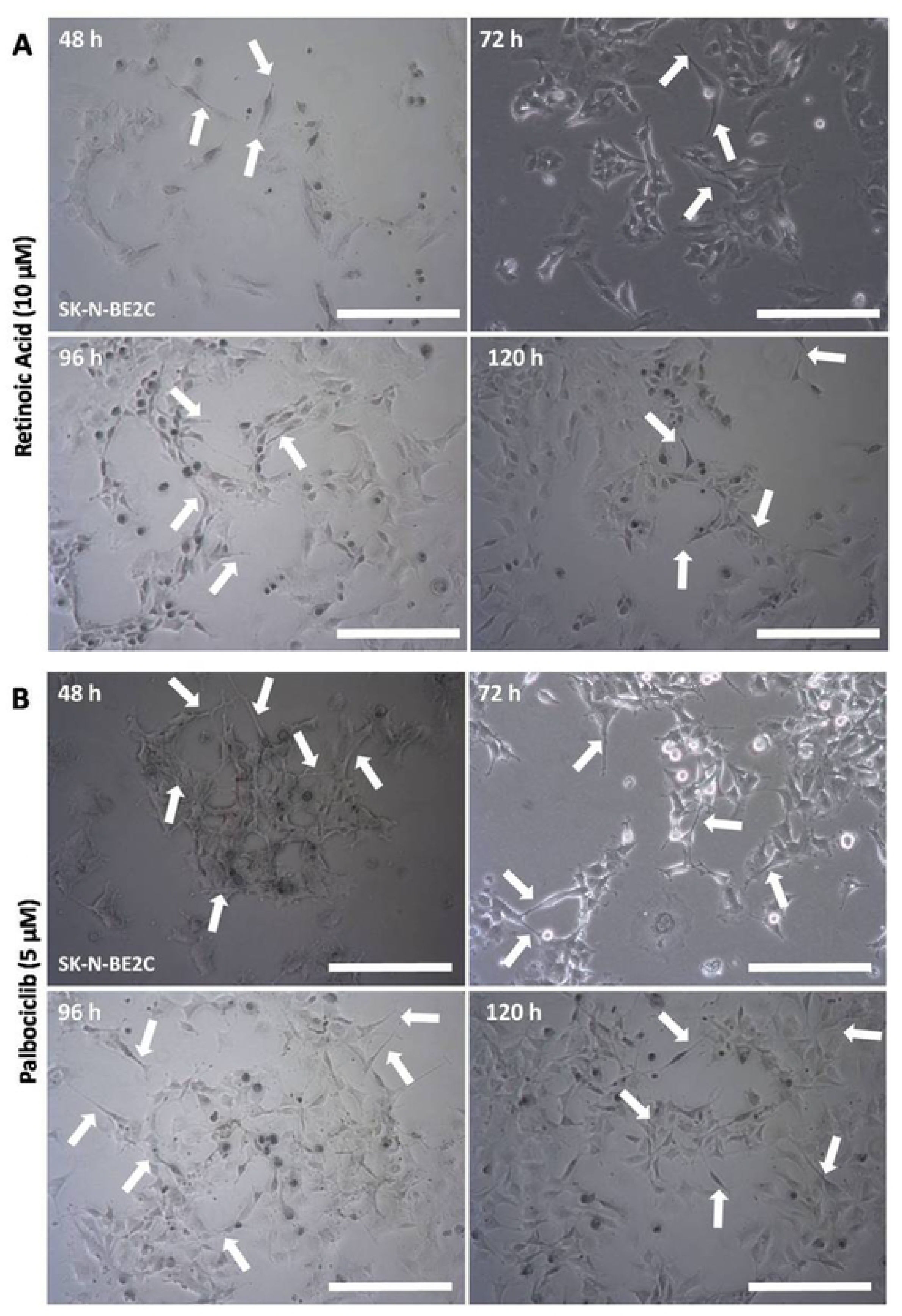
Morphological appearance of replated SK-N-BE2C cells previously treated with RA or Palbociclib. SK-N-BE2C cells previously treated for 72 hours with RA or Palbociclib were passaged and replated in a 75 cm^2^ tissue culture flask in plain cell culture media with no treatment. Phase-contrast images were captured at 48, 72, 96 and 120 hours with Leica microscope x10 objective. Cells continue to produce processes as shown by the white arrows in both (**A**) and (**B**). Cellular processes are much longer in Palbociclib treated cells (**B**) compared to RA treated cells (**A**). This experiment was repeated twice (N=2). Scale bar is 250 µm.

### Cellular differentiation of the NB using Palbociclib

Palbociclib is a known selective inhibitor of cyclin-dependent kinases (CDK4/6) [27,33]. It was the first inhibitor of CDK4/6 approved for cancer treatment. It has been reported to act by inducing differentiation in the aggressive form of breast cancer [28–30]. It was demonstrated in our laboratory that Palbociclib induces neurite formation in SK-N-BE2C cells at 5 µM *in vitro* [10,11]. As such we adopted the same concentration.

SK-N-BE2C cells were plated on glass coverslips, in a 24 well plate. Attached cells were treated for 72 hours with Palbociclib 5 µM at 5% CO_2_ and 37°C. Nuclei were stained with DAPI and images were captured using a confocal microscope. Treatment resulted in neurite formation and was more extensive than seen with RA (**Fig 6**). Our observation was consistent with the previous findings in the lab [10,11]. Further replating of the Palbociclib treated cells in plain culture medium showed that the effect of treatment was sustained as the cells continued to produce neurites even when the drug had been removed (**Fig7 B**).

### Effect of cell differentiation on transendothelial migration of SK-N-BE2C cells

Data in the previous section shows that RA and Palbociclib induced cellular differentiation in SK-N-BE2C cells, as evident by the change in the morphology of the cells, with neurite formation in the treated cells. In addition, the effect of treatment was also shown to be sustained in both RA and Palbociclib treated cells.

To further investigate the effect of cellular differentiation on trans endothelial migration, SK-N-BE2C cells were treated with RA (10 µM) or Palbociclib (5 µM) for 72 hours in a 75 cm^2^ tissue culture flask. The differentiated cells were seeded onto 100% confluent HDMEC monolayers in a 24 well plate, as described previously. Subsequently, live cell imaging was carried out using a cell IQ machine and transmigration was assessed during the 12 hours period. The percentage of non-migrated cells for RA and Palbociclib treated SK-N-BE2C cells was found to be not significantly different to the control (P > 0.05, **Fig 8A**). In addition, a time course analysis shows no difference in the pattern of transmigration of differentiated cells from the undifferentiated cells, for both RA and Palbociclib (**Fig 8B**). Furthermore, the total area under the curve at the end of 12 hours for the control and differentiated cells was almost the same (**Fig 8C**).

Our observations thus show that there was no significant statistical difference in the pattern as well as in the percentage of non-migrated cells between the differentiated and undifferentiated SK-N-BE2C cells for both RA and Palbociclib treatment.

**Fig 8.**
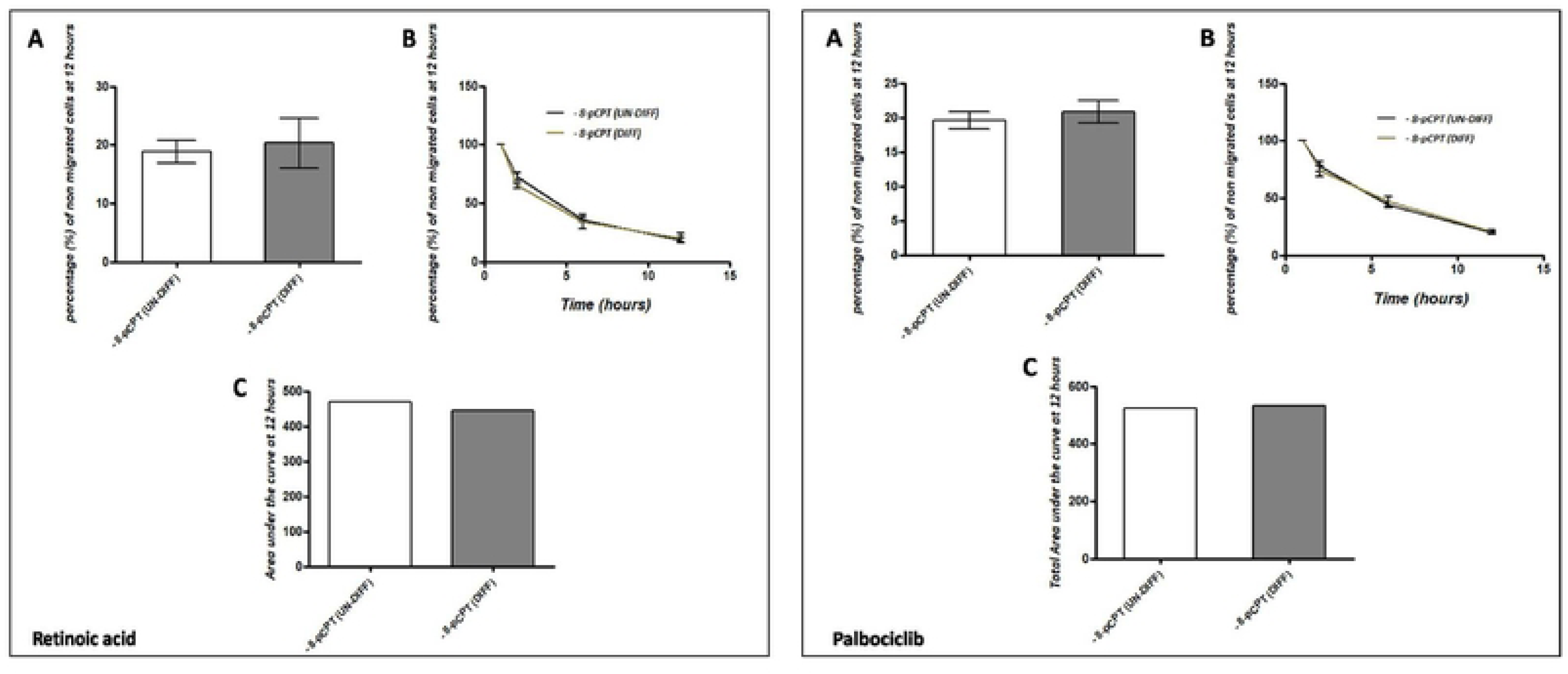
Effect of RA and Palbociclib treatment of SK-N-BE2C cells on trans-endothelial cellular migration. HDMECs were plated on a 0.1% gelatin-coated coverslip in a 24 well plate for 24 hours to generate a confluent monolayer. SK-N-BE2C cells were initially treated with RA (10 µM) and with Palbociclib (5 µM) in separate settings for 72 hours prior to plating on the endothelial cell monolayer. 3 x 10^4^ GFP-labelled differentiated/undifferentiated cells were seeded on a 100% confluent HDMEC monolayer. Serial images were captured at 30-minute intervals using a cell IQ machine. Percentage of non-migrated NB cells were quantified by counting the GFP positive cells at 60 minutes and at the end of 12 hours. Three separate independent experiments were analysed for each of the NB cell lines. Each bar in (**A**) represents the mean ± SEM. Students t-test was used to compare the effect of cellular differentiation on NB cells trans-endothelial migration (P > 0.05). (**B**) Shows a similar time course pattern between RA / Palbociclib differentiated (DIFF) cells and undifferentiated (UN-DIFF) cells. (**C**) Shows the total area under the curve at the end of 12 hours for the non-migrated cells in RA / Palbociclib differentiated as well as the undifferentiated cells. This experiment was repeated three times (N=3).

### Effect of Epac agonist and antagonists on the trans-endothelial cellular migration of differentiated SK-N-BE2C cells

Having established that there was no difference in the percentage and pattern of trans-endothelial cellular migration between differentiated and undifferentiated SK-N-BE2C cells, we then investigated the effect of a single treatment of Epac1 agonist or antagonists on the migration of differentiated cells. Live cell imaging was performed following 72 hours treatment of SK-N-BE2C cells with either 10 µM RA or 5 µM Palbociclib. The Epac antagonists (ESI-09 5 µM and HJC0197 5 µM) did not show any observable difference compared to the controls (**Fig 11A** and **B**, & **Fig 12A** and **B**). However, 8-pCPT treatment in both RA and Palbociclib differentiated cells was found to increase the percentage of non-migrated differentiated cells relative to the control, > 50% in RA differentiated cells and > 90% in Palbociclib differentiated cells (**Fig 9A** and **B**, & **Fig10 A** and **B**).

**Fig 9.**
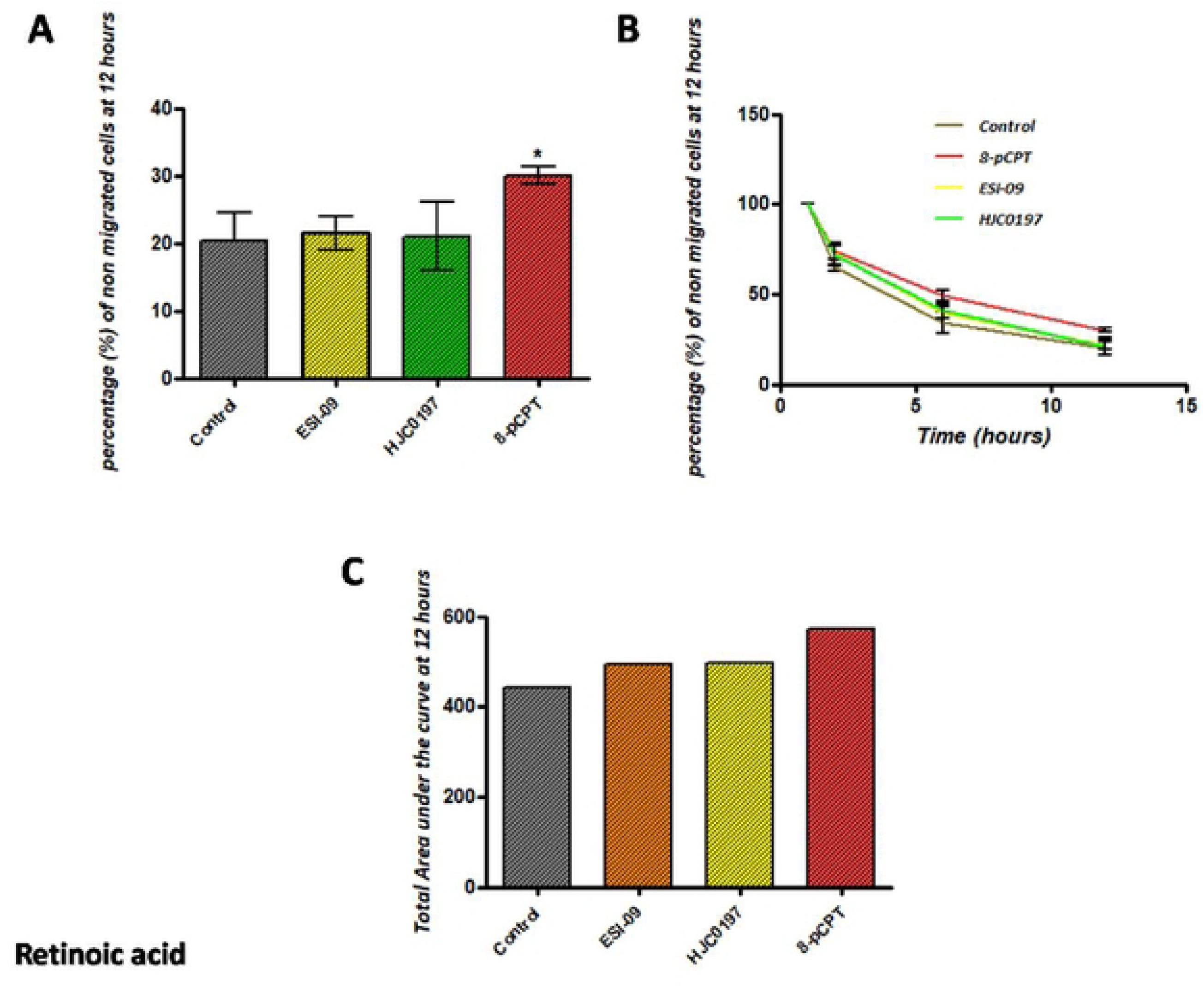
Effect of 8-pCPT, ESI-09 and HJC0197 treatment on trans-endothelial cellular migration of RA differentiated SK-N-BE2C cells. HDMECs were plated on a 0.1% gelatin-coated coverslip in a 24 well plate for 24 hours to generate a confluent monolayer. SK-N-BE2C cells were initially treated with RA (10 µM) for 72 hours prior to plating on the endothelial cell monolayer. 3 x 10^4^ GFP-labelled differentiated / undifferentiated cells were seeded on a 100% confluent HDMEC monolayer. Serial images were captured at 30 minutes intervals using a cell IQ machine. Percentage of non-migrated NB cells were quantified by counting the GFP positive cells at 60 minutes and at the end of 12 hours. Three separate independent experiments were analysed for each condition. Each bar in (**A**) represents the mean ± SEM. One way ANOVA was used to compare the effect of treatment on NB cell trans-endothelial migration. Means comparison was assessed by Tukey post-test (P < 0.05 (*)). (**B**) Shows a similar time course for the antagonists and the control for both differentiated cells and undifferentiated cells. (**C**) Shows the total area under the curve at the end of 12 hours for the non-migrated cells in RA differentiated as well as the undifferentiated cells which was similar for the antagonists and was higher for the 8-pCPT condition. This experiment was repeated three times (N=3).

**Fig 10.**
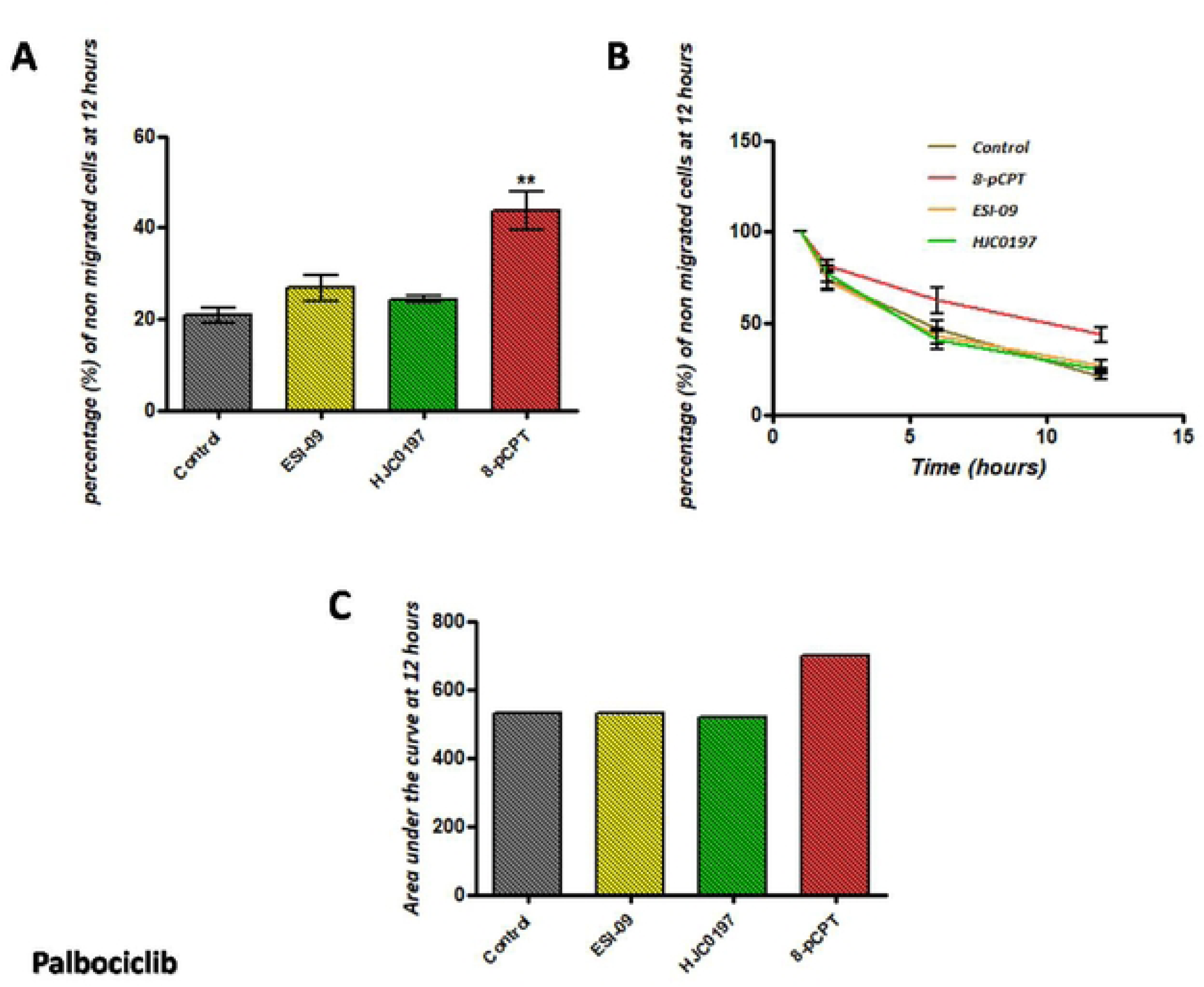
Effect of 8-pCPT, ESI-09 and HJC0197 treatment on trans-endothelial cellular migration of Palbociclib differentiated SK-N-BE2C cells. HDMECs were plated on a 0.1% gelatin-coated coverslip in a 24 well plate for 24 hours to generate a confluent monolayer. SK-N-BE2C cells were initially treated with Palbociclib (5 µM) for 72 hours prior to plating on the endothelial cell monolayer. 3 x 10^4^ GFP-labelled differentiated / undifferentiated cells were seeded on the 100% confluent HDMEC monolayer. Serial images were captured at 30 minutes intervals using a cell IQ machine. Percentage of non-migrated NB cells was quantified by counting the GFP-labelled cells at 60 minutes and at the end of 12 hours. Three separate independent experiments were analysed for each condition. Each bar in (**A**) represents the mean ± SEM. One way ANOVA was used to compare the effect of treatment on NB cells trans-endothelial migration. Means comparison was assessed by Tukey post-test (P < 0.05 (*)). (**B**) Shows a similar time course pattern for the antagonists and the control for both differentiated cells and undifferentiated cells. (**C**) Shows the total area under the curve at the end of 12 hours for the non-migrated cells in Palbociclib differentiated as well as the undifferentiated cells which were similar for the antagonists and higher for the 8-pCPT condition. This experiment was repeated three times (N=3).

Also, time course assessment shows the 8-pCPT treatment delays the transmigration of the differentiated cells compared to the controls for both RA and Palbociclib differentiated cells (**Fig 9B** and **Fig10 B**). The area under the curve of each of the antagonists at the end of 12 hours was found to be similar to the control, which was lower than the 8-pCPT treated condition (**Fig 9C** and **Fig 10C**). Thus 8-pCPT treatment reduces the transmigration of both differentiated and undifferentiated SK-N-BE2C cells.

### Combining Epac agonist with antagonists: Effect on transendothelial migration of differentiated SK-N-BE2C cells

Similar to the single individual treatments for agonist and antagonists, cells were seeded and prepared for trans-endothelial migration assessment. Trans-endothelial cellular migration of the differentiated SK-N-BE2C cells was then compared following combination treatment of 8-pCPT and either of two antagonists (ESI-09 and HJC0197). No significant difference was observed between the groups using agonist and antagonist combination treatments when compared with the control groups (**Fig 11A** & **Fig 12A**) for both RA and Palbociclib differentiated cells. However, we observed a significant increase in the percentage of non-migrated cells in 8-pCPT of >50% in the RA group and >90% in the Palbociclib group (**Fig 11B** and **Fig 12B**). These results were similar to our finding in the earlier study with agonist and antagonist treatment groups.

**Fig 11.**
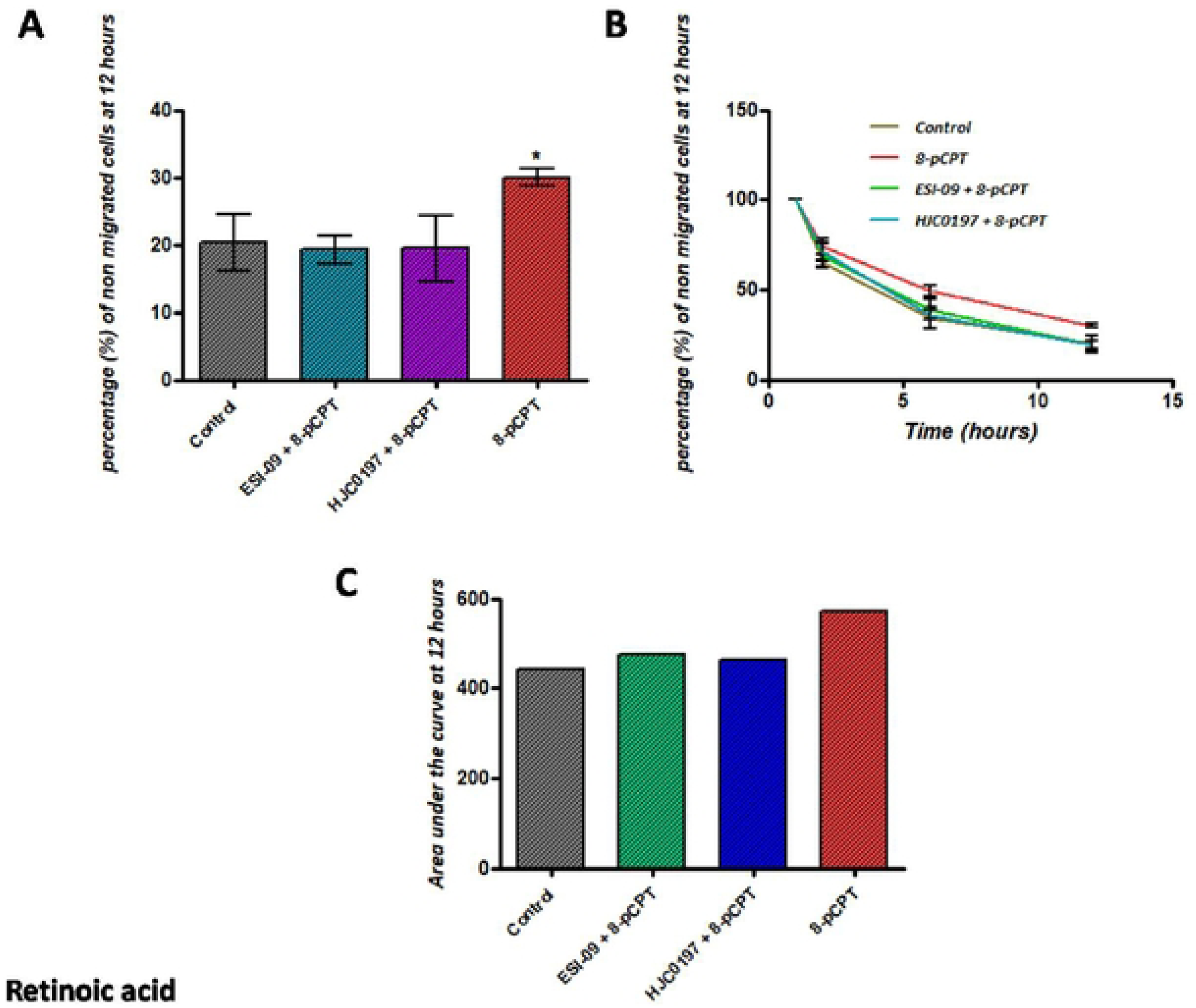
Effect of combination treatment with Epac agonist and antagonists on trans-endothelial cellular migration of RA differentiated SK-N-BE2C cells. HDMECs were plated on a 0.1% gelatin-coated coverslip in 24 well for 24 hours to generate a confluent monolayer. SK-N-BE2C cells were initially treated with RA (10 µM) for 72 hours prior to plating on endothelial cell monolayer. 3 x 10^4^ GFP-labelled differentiated/undifferentiated cells were seeded on 100% confluent HDMEC monolayer. Serial images were captured at 30 minutes interval using cell IQ machine. Percentage of non-migrated NB cells was quantified by counting the GFP-labelled cells at 60 minutes and at the end of 12 hours. Three separate independent experiments were analysed for each condition. Each bar in (**A**) represents the mean ± SEM. One way ANOVA was used to compare the effect of treatment on NB cells trans-endothelial migration. Means comparison was assessed by Tukey post-test. P < 0.05 (*). (**B**) Shows a similar time course pattern between the antagonists and the control for both differentiated cells and undifferentiated cells. (**C**) Shows the total area under the curve at the end of 12 hours for the non-migrated cells in RA differentiated as well as the undifferentiated cells which were similar for the antagonists and higher for the 8-pCPT condition. This experiment was repeated three times (N=3).

**Fig 12.**
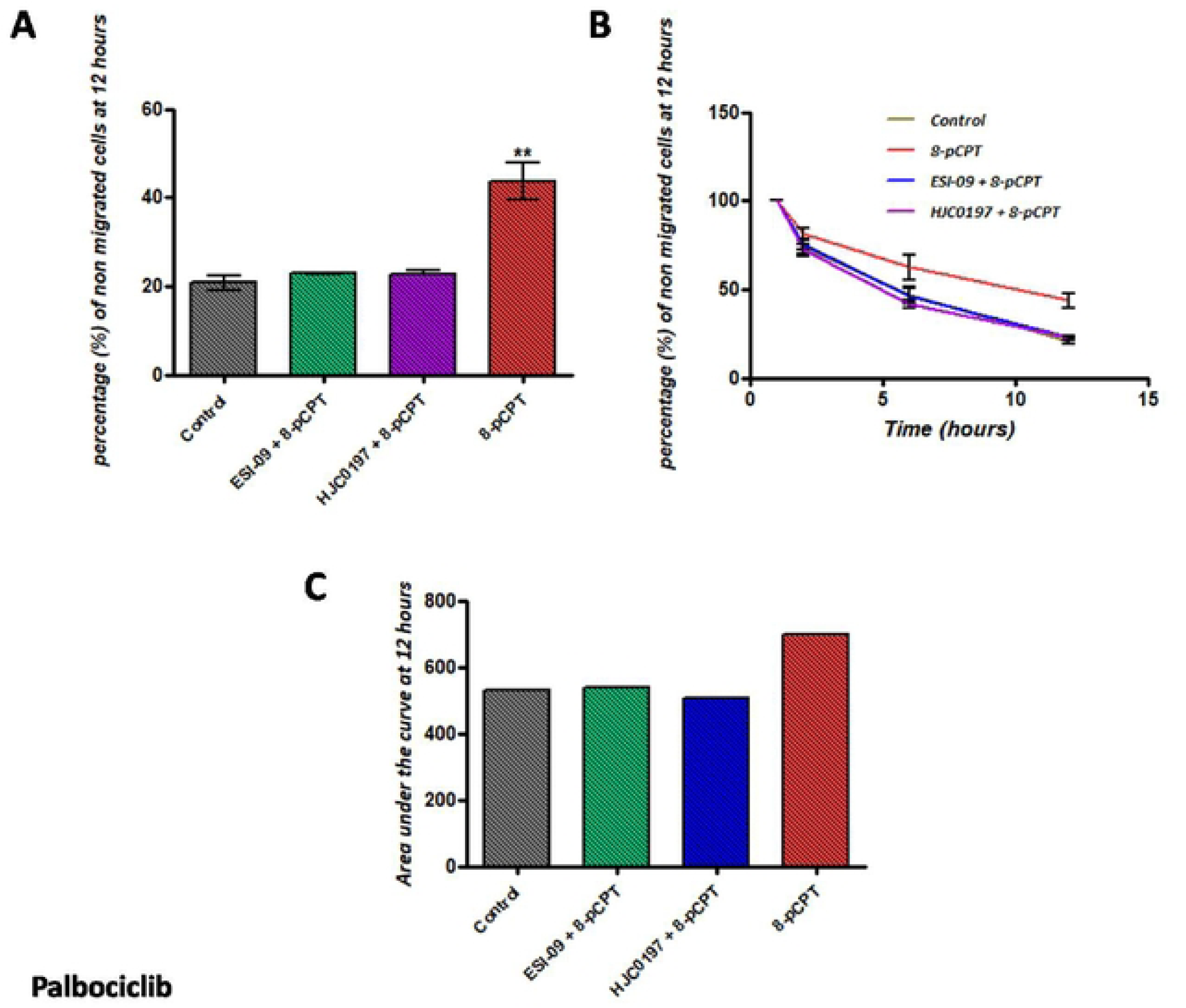
Effect of combination treatment with Epac agonist and antagonists on trans-endothelial cellular migration of RA differentiated SK-N-BE2C cells. HDMECs were plated on a 0.1% gelatin-coated coverslip in 24 well for 24 hours to generate a confluent monolayer HDMEC cells were plated on a 0.1% gelatin-coated coverslip in 24 well for 24 hours to generate a confluent monolayer. SK-N-BE2C cells were initially treated with Palbociclib (5 µM) for 72 hours prior to plating on endothelial cell monolayer. 3 x 10^4^ GFP-labelled differentiated / undifferentiated cells were seeded on 100% confluent HDMEC monolayer. Serial images were captured at 30 minutes interval using cell IQ machine. Percentage of non-migrated NB cells was quantified by counting the GFP-labelled cells at 60 minutes and at the end of 12 hours. Three separate independent experiments were analysed for each condition. Each bar in (**A**) represents the mean ± SEM. One way ANOVA was used to compare the effect of treatment on NB cells trans-endothelial migration. Means comparison was assessed by Tukey post-test. P < 0.01 (**). (**B**) Shows a similar time course pattern between the antagonists and the control for both differentiated cells and undifferentiated cells. (**C**) Shows the total area under the curve at the end of 12 hours for the non-migrated cells in Palbociclib differentiated as well as the undifferentiated cells which were similar for the antagonists and higher for the 8-pCPT condition. This experiment was repeated three times (N=3).

As anticipated the total area under the curve at the end of 12 hours for the agonist treatment in both RA and Palbociclib groups of differentiated cells was found to be higher compared to the agonist/antagonists treatment combination (**Fig 11C** and **Fig 12C**). The time course analysis shows that 8-pCPT treatment delays the transmigration of the differentiated cells compared to the controls for both RA and Palbociclib differentiated cells (**Fig 11B** and **Fig 12B**).

## Discussion

Metastasis is a highly complex event in neuroblastoma as well as in other cancers. The overall survival in high-risk patients is <50% even with treatment, signifying the need for further research towards the development of new additional therapies [34–36]. Here we set up a good, reliable as well as reproducible co-culture model (**Fig 1**) with the aid of Cell IQ machine to assess the effect of Epac agonist on trans-endothelial migration of neuroblastoma cells, by means of co-culturing neuroblastoma and endothelial cells *in vitro*. Co-culture treatment with 8-pCPT alone in both SK-N-BE2C and SK-N-AS showed a reduction in the number of cells traversing the endothelial monolayer, as depicted by an increase in the percentage of the non-migrated NB cells. The increase in percentage of non-migrated NB cells could be associated with strengthening of the VE-cadherin adherens junction caused through the activation of Epac [12,15,17,18]. Wittchen and colleagues have further demonstrated that in addition to enhancing the endothelial barrier function, activation of the Epac signalling pathway also inhibits trans-endothelial migration of differentiated HL60 cells through elevating the activity of Rap1 within the endothelial cells [37]. Interestingly we observed the antagonists to perfectly inhibit the activation of Epac 1 by the agonist, as expected (20,38). These findings thus suggest the potential therapeutic role of 8-pCPT in the control of metastatic cancer.

RA [31,32] as well as Palbociclib [34,39] differentiated SK-N-BE2C cells (**Fig 6** and 7). Differentiated cells co-cultured with HDEMCs, upon 8-pCPT treatment, showed a two-fold increase in the percentage of retained cells for the Palbociclib differentiated compared to the RA differentiated cells (**Fig 9** and 10). The RA differentiated cells behaved in a similar pattern to the undifferentiated cells. One possible explanation could be that Palbociclib differentiation may cause cells to join one another, forming a cellular sheath that is unable to easily transmigrate, contrary to the RA differentiated cells. Additionally, it could be that 8-pCPT is activating some cascade of events in Palbociclib differentiated cells resulting in inhibition of transmigration and this has to be further investigated. By and large, Palbociclib differentiated cells demonstrated larger fold reduction of transmigration.

In conclusion we established that tightening of the endothelial barrier achieved sequel to Epac activation in HDMECs reduced the trans-endothelial migration of the NB (SK-N-BE2C and SK-N-AS) cells, which potentially may serve as a therapeutic mechanism for the Epac pathway in the treatment of the high-risk neuroblastoma patients as they often developed the refractory or recurrent osteomedullary form of NB disease.

## Acknowledgements

RI was supported by grants from Tertiary Education Trust Fund (tEtfund), Bayero University Kano in Nigeria. This work also was funded by Alder Hey Oncology fund (DM and JQ).

## Supporting information

**S1 Fig. Effect of combination treatment with Epac agonist and antagonists on SK-N-BE2C and SK-N-AS migration.**. Comparison of SK-N-BE2C and SK-N-AS cells in wound closure assay. The cells were treated with either forskolin (10 µM) alone or with forskolin (10 µM) and Epac antagonist (HJC0197 (5 µM) or PKI (5 µM)). No difference was observed between the forskolin-treated and untreated SK-N-BE2C and SK-N-AS cells. Wound treated with PKA inhibitor and Epac antagonists did not change the speed of migration. Each plot in graphs for SK-N-BE2C and SK-N-AS represents the mean ±SEM. This experiment was repeated three times (N=3).

**Fig S2. Effect of treatment with 8-pCPT, ESI-09 and HJC0197on SK-N-BE2C and SK-N-AS proliferation.** SK-N-BE2C and SK-N-AS cells were seeded into a flat bottom 96 well plate for 24 hours. Cells were then treated with 8-pCPT (10 µM), ESI-09 (5 µM) and HJC0197 (5 µM) for 24 hours and MTT absorbance were measured. 8-pCPT, as well as ESI-09 and HJC0197 treatment, did not cause any clear change in the cell number in both cell lines compared to the control (MED). Each bar represents the mean ± SEM. One way ANOVA was used to compare the effect of treatment on NB cells proliferation. Means comparison was assessed by Tukey post-test. P > 0.05. This experiment was repeated three times (N=3).

